# Heat shock factor 5 is conserved in vertebrates and essential for spermatogenesis in zebrafish

**DOI:** 10.1101/254615

**Authors:** Jolly M. Saju, Mohammad Sorowar Hossain, Woei Chang Liew, Ajay Pradhan, Natascha May Thevasagayam, Amit Anand, Per-Erik Olsson, László Orbán

**Author notes:** Addresses for correspondence:* László Orbán, Department of Animal Sciences, Georgikon Faculty, Pannon University, Keszthely, Hungary; Per-Erik Olsson, Biology, The Life Science Center, School of Science and Technology, Örebro University, Örebro, Sweden; Amit Anand, Bioimaging and Biocomputing, Temasek Life Sciences Laboratory, 1 Research Link, The NUS, Singapore 117604.

## Abstract

Heat shock factors (Hsfs) are transcription factors that regulate response to heat shock and to variety of other environmental and physiological stimuli. Four HSFs (HSF1-4) known in vertebrates till date, perform a wide variety of functions from mediating heat shock response to development and gametogenesis. Here, we describe a new yet conserved member of HSF family, Hsf5, which likely exclusively functions for spermatogenesis. The *hsf5* is predominantly expressed in developing testicular tissues, in comparison to wider expression reported for other HSFs. HSF5 loss causes male sterility due to drastically reduced sperm count, and severe abnormalities in remaining few spermatozoa. While hsf5 mutant female did not show any abnormality. We show that Hsf5 is required for progression through meiotic prophase 1 during spermatogenesis. The *hsf5* mutants indeed show misregulation of a substantial number of genes regulating cell cycle, DNA-damage repair, apoptosis and cytoskeleton proteins. We also show that Hsf5 physically binds to majority of these differentially expressed genes, suggesting its direct role in regulating the expression of many genes important for spermatogenesis.

## Introduction

Heat shock transcription factors (Hsfs) are involved in the differentiation, development, reproduction and stress-induced adaptation by regulating temperature-controlled heat shock protein (*hsp*) genes and other, non-hsp genes as well ((1, 2); (for the complete list of genes analyzed in the study see (**Supplementary Table 1A**). Hsps are molecular chaperones that maintain the cellular homeostasis and promote survival (3, 4). Multiple Hsfs in plants and vertebrates appear to mediate a wide array of responses to versatile forms of physiological and environmental stimuli (5).

Four members of the Hsf family have been identified and characterized in vertebrates prior to this study (reviews: (1, 6, 7)). Three of them (Hsf1, Hsf2 and Hsf4) are conserved in all vertebrate groups, whereas the fourth (Hsf3) is found in mouse, birds, lizards and frogs. The canonical Hsf1 and Hsf3 have been shown to mediate stress-induced Hsp expression in response to various environmental stressors such as elevated temperatures and cadmium sulfate, or exposures to proteosome inhibitors (8-10). In the case of Hsf1, the stress responses are generally manifested by the ability of this transcription factor to exhibit inducible DNA binding activity, nuclear localization and oligomerization (11). Mouse cells with inactivated Hsf1 have been shown to be unresponsive to stress-induced *Hsp* gene expression (12). Hsf1-knockout mice were found to show enhanced effects upon treatments with cadmium (13) and bacterial endotoxins (12).

On the other hand, Hsf2 and Hsf4 were not stress-responsive; their expression varied strongly during differentiation and development. In Hsf2-knockout mice, microarray-based transcriptomic analysis of embryos and testis – in comparison to controls - could not detect any differentially expressed (DE) genes from the HSF/chaperone family (14). Hsf4 was found to be involved in lens development in the rat (15). Immunohistochemical analysis of human and mouse testis sections with an anti-Hsf5 antibody showed highest staining intensity in spermatocytes followed by spermatids (16). In vertebrates, HSFs play a crucial role in the maintenance of the reproductive function, mainly during spermatogenesis (1). In Drosophila, there is only a single Hsf, which regulates heat-shock induced gene expression, and it is also essential for larval development and oogenesis (17).

Zebrafish (*Danio rerio, Cyprinidae*) is an important vertebrate model organism, which has helped to answer important biological questions related to development, genetics and diseases (18-20). This small freshwater species offers several advantages over other models due to its small size, short generation time, transparent embryonic development and availability of relatively large number of eggs on a weekly basis. Zebrafish is also suitable for high throughput experiments and large-scale mutagenesis for genetic studies. Prior to this study, three heat shock factors, namely Hsf1, Hsf2 and Hsf4 have been isolated and characterized from zebrafish (14, 21, 22).

In order to understand zebrafish reproduction at the molecular level, others and we have performed comparative expression analyses of male vs. female gonads at several different developmental stages from 21 days post-fertilization (dpf) to adults (23-25). Array-based transcriptomic studies performed in our lab earlier on adult gonads identified a number of novel genes with gonad-specific (or gonad-enhanced) expression (23, 25). One of these gonad-enhanced novel genes was *heat shock factor 5 (hsf5)*, which is a new member of the Hsf family. We cloned the full-length cDNA of *hsf5* from zebrafish and characterized its expression in embryos, developing gonads and adult tissues. In adult zebrafish, the gene is expressed in several adult organs, with the testis showing higher transcript levels than other organs, including the ovary, and a short transcript variant specific to testis. We have created zebrafish mutants of *hsf5* by CRISPR/Cas9 technology (26), and used them for basic functional characterization of the gene product. Testes of adult *hsf5*^-/-^ *males* were primarily dominated by spermatocytes with very few spermatozoa remaining. A detailed analysis of mutants revealed that *hsf5* loss-of-function results in failure of progression of meiotic prophase 1, through misregulation of cell cycle, meiosis and DNA repair. The head size of mutant spermatozoa has increased compared to control and many had distorted or missing flagella. While all *hsf5*^-/-^ males were infertile, *hsf5*^-/-^ *and hsf5*^+/-^ females as well as *hsf5*^+/-^ males exhibited normal gonadal appearance and fertility. To the best of our knowledge, this is the first functional characterization of Hsf5 from any of the vertebrate species and the first molecular insight for its role in zebrafish spermatogenesis.

## Results

### Identification and cloning of Hsf5, a new heat shock transcription factor

We identified *hsf5* initially as an EST with enhanced expression in zebrafish testis from a normalized adult testis cDNA library (23). Subsequent RACE analysis and sequence comparison with other ESTs led to identification of full-length 1.7 kb transcript (FJ969446, NM_001089476).

Mapping of the full-length cDNA to the latest genome assembly (Genome Reference Consortium, GRCz11; released in May 2017) revealed that the genomic locus consisted of six exons spanning a region of over 20 kb on Chromosome 5 (3,094,980 - 3,118,313 bp). Bioinformatic analysis of the predicted protein revealed that the N-terminal region (between the 19^th^ and 119^th^ amino acids) of the protein contains a helix-turn-helix DNA binding domain (DBD) that is the most conserved functional domain in HSFs across vertebrates **(Fig. 1A)**. The DBD of the new gene showed 37-39% identity with those of other zebrafish Hsfs **(Supplementary Table 1D)**.

**Fig. 1:**
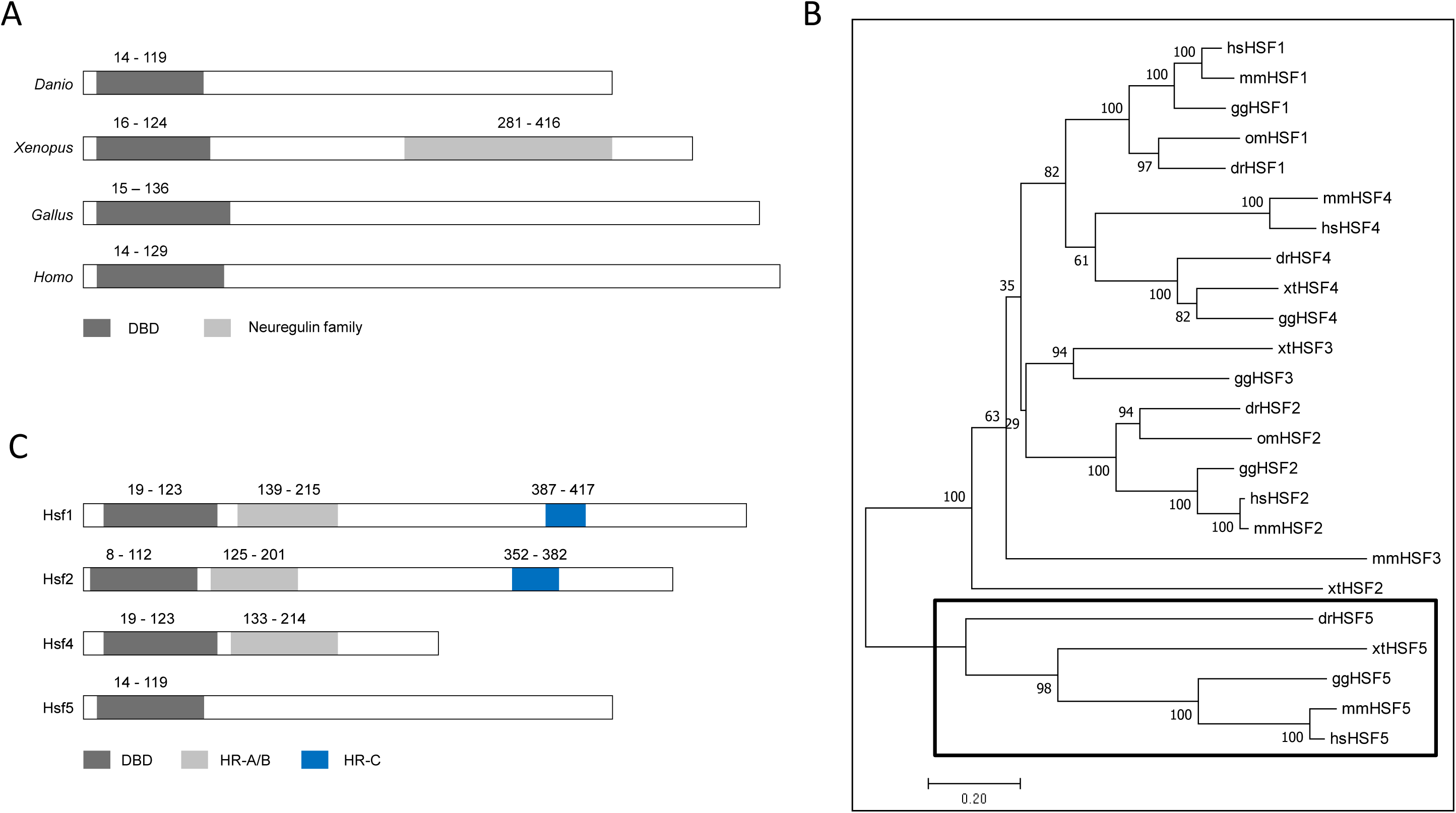
The new heat shock factor (Hsf5) isolated from zebrafish can be found in members of all major vertebrate groups and lacks HR-A/B and HR-C domains. A: Comparative view of the domain architecture of four vertebrate Hsf5s: *Danio* - zebrafish, *Xenopus* - frog, *Gallus* - chicken and *Homo* - human. DBD - DNA-binding domain; B: Phylogenetic analysis of vertebrate HSFs showed that all orthologs of the newly described zebrafish gene are located on a separate branch clearly apart from the rest of the other family members (HSF1-4). The tree was constructed with the neighbour-joining method based on the amino acid sequences; the confidence in phylogeny was assessed by bootstrap re-sampling of the data (1000X). Labels: hs - *Homo sapiens*, mm - *Mus musculus*, gg - *Gallus gallus*, xt - *Xenopus tropicalis*, om - *Oncorhynchus mykiss*, and dr -*Danio rerio* (GenBank IDs of all sequences used are shown in **S1A Table**). C: Schematic representation of the four members of zebrafish heat shock factor (Hsf) protein family for comparing the relative location of specific domains. HR-A/B - heptad repeat (A/B); HR-C - heptad repeat (C).

Orthologs were identified in human (NP_001073908.2), mouse (NP_001038992.1), chicken (XP_003642431.2), python (XP_007429104), Xenopus (NP_001107312.1) and guppy (XP_008426339), thus we propose that this protein is likely to be conserved in most vertebrates. On the phylogenetic tree, all previously described Hsfs were grouped into four separate clusters (Hsf1-4), whereas the new gene and its presumed vertebrate orthologs have formed a new branch **(Fig. 1B)**. These data confirm that this is the latest, novel member of the heat shock factor family and is classified as Hsf5.

In addition to the DBD domain, we could not find other domains in zebrafish Hsf5; the Neuregulin domain found in Xenopus Hsf5, and the heptad repeat A/B and heptad repeat C (HR-C) domains present in the other Hsfs were not present **(Fig. 1C)**. Zebrafish Hsf5 shares structural similarities with its orthologs from frog, chicken and human HSF5, including the lack of HR-A/B and HR-C domain. This indicates that Hsf5 monomers and dimers may not be able to form trimers, bind to a typical heat shock element or induce heat-shock response. We examined experimentally whether *hsf5* expression is altered upon heat shock as observed for many other heat shock response proteins. The expression level of *hsf5* did not increase upon heat shock in adult testis, whereas transcript levels of *hsp70* showed a significant up-regulation, suggesting that Hsf5 may not be directly involved in heat shock response **(Supplementary Fig 2B-C)**. The total number of heat shock factors in zebrafish now stands at four: Hsf1 (27), Hsf2 (22), Hsf4 (21) and Hsf5 (this publication).

### Hsf5 shows sexually dimorphic expression

The *hsf5* transcript was maternally deposited in zebrafish oocytes and the first sign of its zygotic expression was observed at mid-blastula transition stage **(Supplementary Fig 3A)**. In adults, *hsf5* expression was significantly higher in the testis than other organs, including the ovary **(Supplementary Fig 3B)**.

The male-specific expression pattern of *hsf5* was established during the development. In comparison to two early testicular markers, *amh* and *nr5a1a*, and ovarian marker *cyp19a1a*, which showed up-regulated expression in respective gonads from 30 dpf (28, 29), *hsf5* expression remained low in developing gonads from both sexes until 35 dpf **(Supplementary Fig 4)**. The *hsf5* expression levels increased significantly in testis from 35 dpf, while its expression at low levels continued in all other tissues, including ovaries, beyond that timepoint **(Supplementary Fig 4D, 5A)**. The above expression profile of *hsf5* suggests that the protein likely functions in the downstream processes of the gonadal transformation towards the maturation of testis.

RT-PCR and sequencing of full-length *hsf5* revealed the presence of an additional, shorter transcript variant *(hsf5_tv2)* in testis, but not in the ovary **(Supplementary Fig 5A)**. The expression level of the short variant was substantially lower than that of the long one *(hsf5_tv1)*. Sequence comparison revealed that the shorter variant lacked the third exon, resulting in the reduction of coding region to 1008 bp from 1251 bp of the long one **(Supplementary Fig 5B)**.

### Generation of Hsf5 mutants in zebrafish using CRISPR-Cas9

First, we tried to ablate Hsf5 function by MO-based knockdown; when tested in embryos, the MO-injected samples did not show any specific phenotype different from mock-injected controls **(Supplementary material S12)**. Similarly, none of the heat or chemical treatments on testicular explants yielded any significant change in the expression level of *hsf5* **(Supplementary material S12)**. Therefore, we decided to generate loss-of-function mutants by CRISPR-Cas9 technology.

Altogether, 16 founders have been validated through sequencing of the targeted exon (exon #2). Ten of them were outcrossed with WT partners to generate F1 offspring **(Supplementary Fig 6)** and heterozygous mutants were identified by fluorescent PCR with primers flanking the deletion site (see Materials and Methods). Heterozygous mutant siblings were crossed to generate the F2 progenies and a typical Mendelian segregation was observed. Three mutant lines were selected for downstream analysis and were named as Hsf5^sg40^ Hsf5^sg41^ and Hsf5^sg42^ as per ZFIN zebrafish nomenclature guidelines. Each line carried a different mutation; a 5bp and 7bp deletion and a 25 bp insertion, respectively, at the expected site in exon #2 coding for DNA binding domain(DBD). At protein level all three selected mutations in *hsf5* gene caused a frameshift leading to a premature stop codon, generating truncated proteins **(Fig. 2A-B)**. Since these mutations result in early termination of hsf5, in the DNA binding domain, we speculate that if the truncated Hsf5 is expressed in mutants, its function will be drastically affected. From here onwards, we refer to all these mutant alleles collectively as *hsf5*^-/-^.

**Fig. 2:**
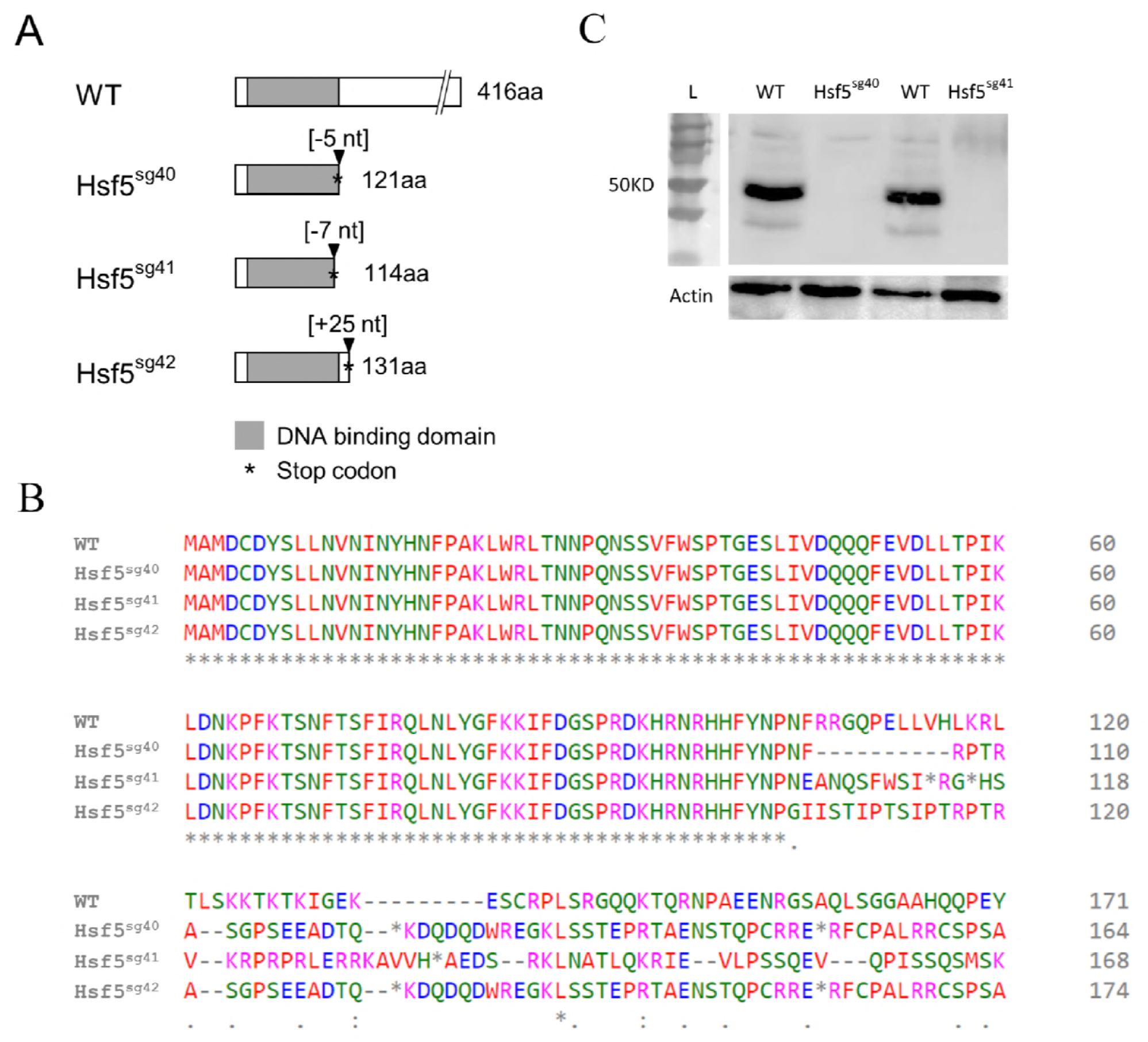
Three hsf5^-/-^ mutant lines were generated by CRISPR/Cas9-based knockout. A: Schematic representation of three mutant lines. Hsf5^sg40^ has a deletion of five bases and it introduces a stop codon after 121 amino acids. Hsf5 ^sg41^ and Hsf5^sg42^ has a deletion of 7 bases and addition of 25 bases and introduces the stop codon after 114 and 131 amino acids respectively and all three mutations are frame shift type resulting in truncated protein. B: Predicted amino acid sequence from three mutant lines. **C:** Western blot confirms the efficiency of the CRISPR/Cas9 treatment. Knockdown of Hsf5 resulted in severe reduction of the protein level compared to WT. Adult testis from Hsf5^sg40^ and Hsf5^sg41^ and WT were used. Actin was used as an internal control.

Homozygous mutants carrying a 7bp deletion (labeled as Hsf5^sg41^ **in Fig. 2A**) were used for Western blot, histology, SEM and RNAseq studies. Western blot analysis with anti-Hsf5 antibody, in Hsf5^sg40^ and Hsf5^sg41^ could not detect any Hsf5 protein confirming the loss of functional, full length Hsf5 **(Fig. 2C)**. To examine the role of *hsf5* in heat shock response, we performed heat treatment on the hsf5 mutants. Expression of *hsp70* in the hsf5 mutants was comparable to that in the WT heat exposed individuals **(Supplementary Fig 2B)**. *hsp70* induction in *hsf5* mutants suggests that Hsf5 is unlikely to play a direct role in heat shock response **(Supplementary Fig 2C)**.

#### 3.4 Hsf5 is predominantly expressed in spermatocytes

Immunostaining revealed abundant Hsf5 expression in testis (**Fig. 3A**). Hsf5 expression was predominantly observed in spermatogonia and primary spermatocytes, whereas its expression in spermatids or spermatozoa was significantly low **(Fig. 3B)**. We compared the localization of Hsf5 with that of a known primary spermatocyte marker, Sycp3 and found that both share similar localization patterns in spermatocytes **(Fig. 3C-D)**. Expectedly, immunostaining and western blotting showed that Hsf5 expression in the ovary was much lower **(Fig. 3F, Supplementary Fig 2A)** than that in the testis. A closer examination of Hsf5 localization in testis revealed that Hsf5 is also localized as foci in the nucleus, while bulk of Hsf5 signal was outside of nucleus **(Fig. 3 G-H)**.

**Fig. 3:**
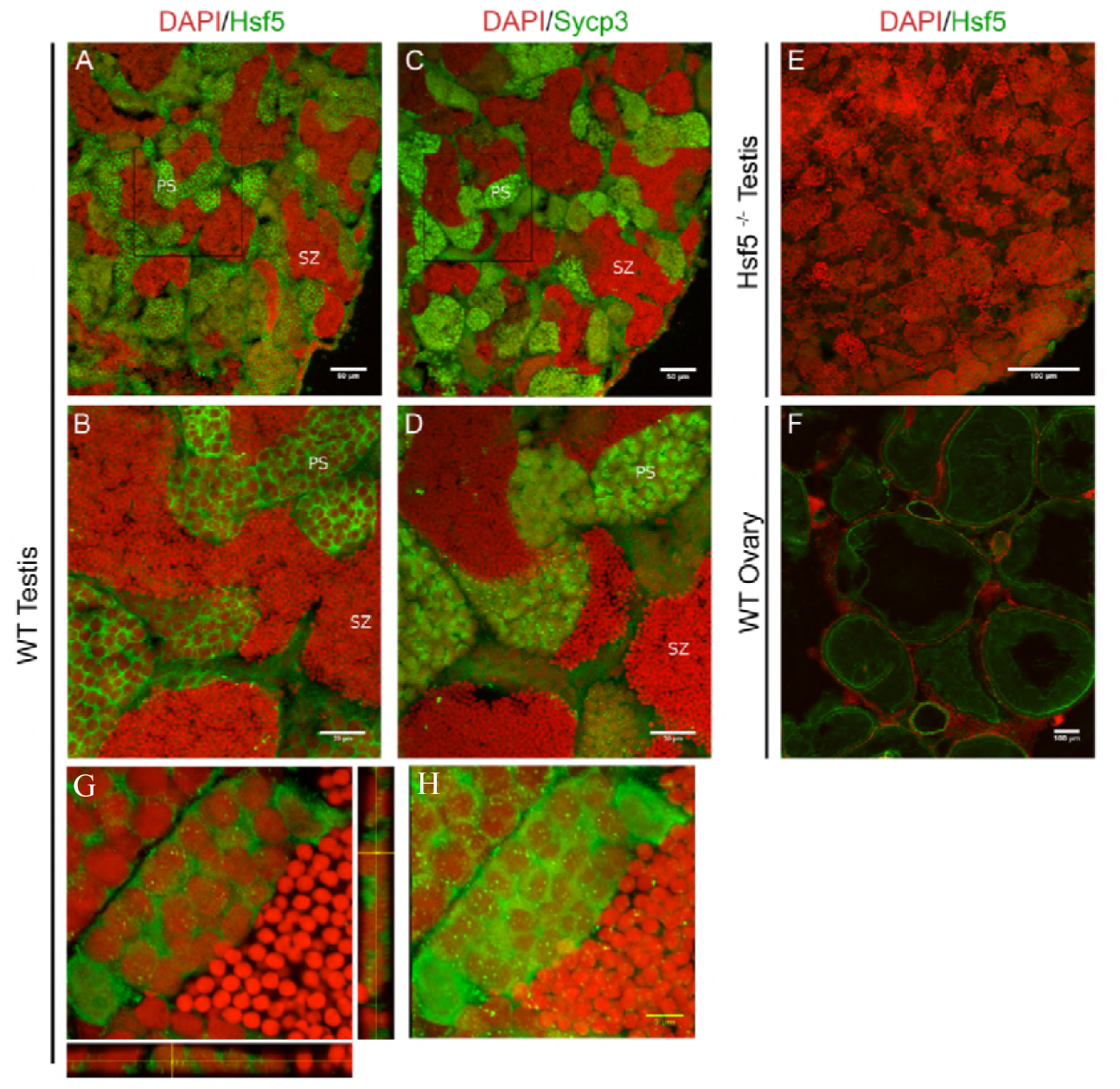
Hsf5 protein showed similar localization pattern to a known spermatocyte marker Sycp3. A-D. Immunohistochemistry was performed on sequential sections from the testis of WT adult zebrafish for Hsf5 and Sycp3 and counterstained with DAPI. **E-F:** Hsf5 expression was undetectable Hsf5^-/-^ testis (E) and was drastically low in WT ovary (F). **G:** Localization of Hsf5 protein (green) as granule like structures in the nucleus of primary spermatocytes, orthogonal views are presented to emphasize presence of foci in nucleus (red). H: Maximum-intensity Z-projection of images. Scale bars - A-D: 50 μm, E: 400 μm, F: 100 μm, G and H: 5 μm

### Hsf5 is required for proper spermatogenesis and fertility in males

When crossed with WT partners, homozygous *(hsf5*^-/-^) and heterozygous *(hsf5*^+/-^) females as well as heterozygous males produced viable offspring, whereas homozygous mutant males failed to generate any viable offspring even after multiple trials. When embryos from the latter crosses were examined under light microscope, they exhibited a substantial delay in their early development and eventual lethality before the age of one dpf **(Supplementary Fig 7)**.

Infertility in homozygous male mutants prompted us to perform a histological comparison of wild type and *hsf5*^-/-^ mutant testes. Expectedly, spermatids and well-developed lumina filled with spermatozoa could be observed in the wild type **(Fig. 4A)**, whereas the *hsf5*^-/-^ mutant testis exhibited a drastic loss of spermatozoa **(Fig. 4B)**. A quantitative comparison of cell types between the mutant and WT testes revealed a significant increase in the number of primary spermatocytes and drastic reduction of spermatozoa in *hsf5*^-/-^ mutant testis **(Fig. 4C)**, whereas the number of spermatogonia was comparable. Analysis of *hsf5*^-/-^semen smear under light microscope showed a drastic reduction in sperm counts in the hsf5 mutants compared to that in the WT (**Fig. 4D**). The *hsf5* mutants also exhibited lower sperm motility in mutants than those of wild types (data not shown). Briefly, spermatogonia are the largest germ cells with large nucleus and poorly condensed chromatin, primary spermatocytes are characterised by the coarse chromatin strands and bouquet/umbrella configuration of chromosomes, and spermatid/spermatozoa are dark round nuclei with minimal or no cytoplasm counted together as one group **(Fig. 4E-G)**.

**Fig. 4:**
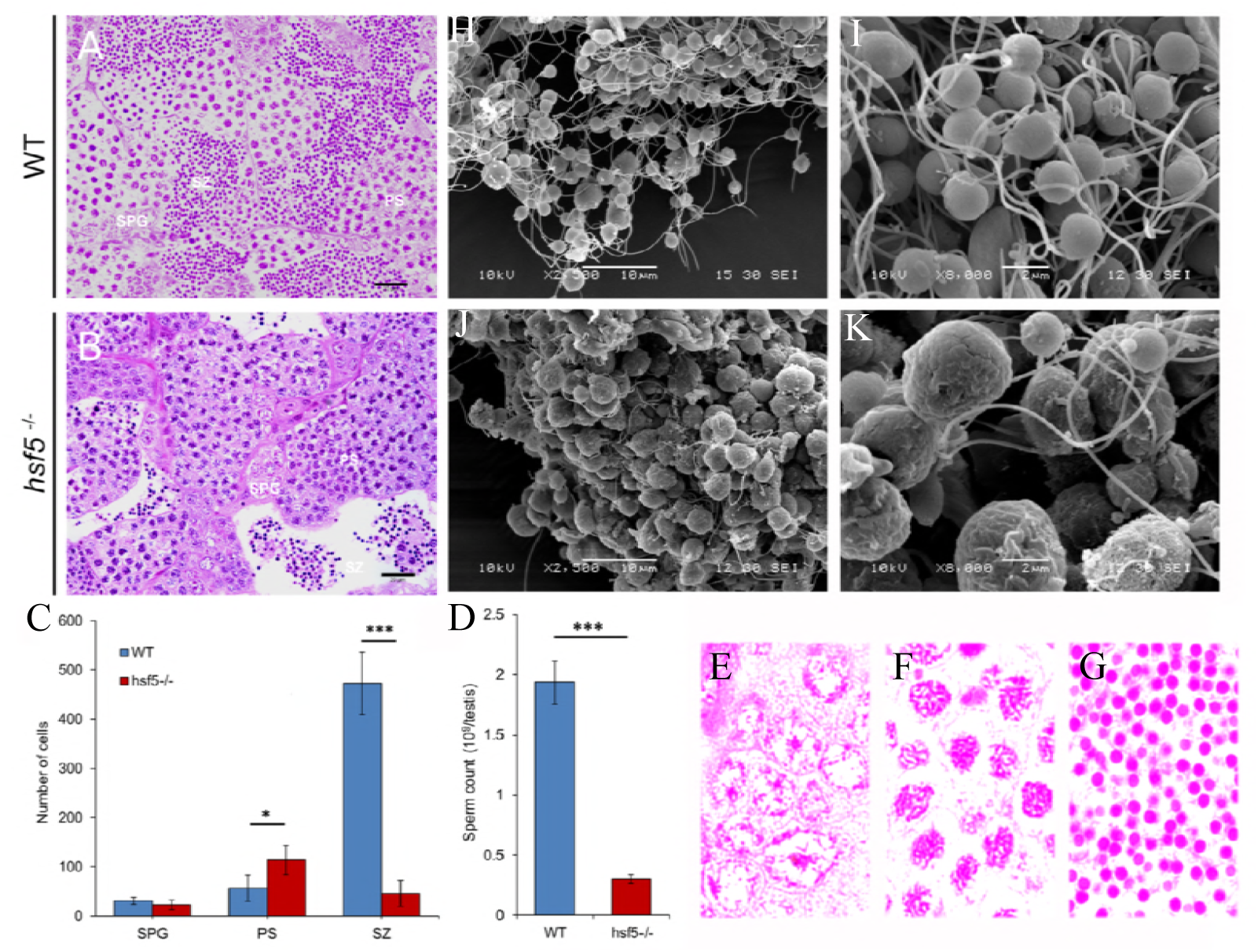
Histological and EM-based analysis of testes and spermatozoa revealed severe reduction and defective shape of spermatozoa in *hsf5*^-/-^mutant compared to WT. Histological analysis showed lumen filled with spermatozoa (SZ), are visible in the wild type (A), and fewer in the mutant (B). In the wild type (A), meiotic prophase stages were visible in few cells, whereas in the mutant (B), an over representation of primary spermatocyte (PS) was observed. C: Quantification of germ cells at different stages in WT and *hsf5*^-/-^ testis. D: The sperm count determined by hemocytometer showed a severe reduction in the number of spermatozoa in *hsf5*^-/-^ compared to WT. E-G: Representative germ cell stages, spermatogonia (spg), primary spermatocyte (ps) and spermatozoa (sz). H-K: Scanning electron micrograph (SEM) of WT (H, I) and *hsf5*^-/-^ spermatozoa (J, K). Spermatozoa of the *hsf5*^-/-^ mutant were clumped together with enlarged head and defective and shorter flagellum.

**Fig. 5:** Transmission electron micrograph (TEM) showing abnormal head axoneme and filament of *hsf5*^-/-^ mutant spermatozoa. A-D: Sperm head from WT (A) and *hsf5*^-/-^ (B-D) showed dislocated cytoplasmic membrane and intense vacuolization in mutants compared to WT. E-F: Cross-section through the central region of a wild type (E) and *hsf5*^-/-^ mutant (F-H) sperm flagella showing the normal 9+2 organisation of the axoneme in WT and variability of the axoneme structure in *hsf5*^-/-^ with defects in central and peripheral duplets symmetry. I-J: Longitudinal section of wild type (I) and *hsf5*^-/-^ mutant (J) flagella showing the defects of microtubules. K: Defects in head and microtubule arrangements in axonemes of wild type (WT) and *hsf5*^-/-^ spermatozoa were quantified and are shown as percentages for each of the categories represented in the table

Next, we examined the sperm morphology in detail, using scanning electron microscope. The results revealed a high proportion of grossly enlarged sperm heads in the *hsf5*^-/-^ mutant compared to controls **(Fig. 4H-K)**. Mutant sperm cells appeared clustered together with crenate arrangement of cytoplasmic membrane. The flagella were either too short or absent from many of the mutant sperm cells compared to wild type **(Fig. 4H-K)**.

A detailed analysis of *hsf5*^-/-^ spermatozoa with transmission electron microscopy revealed irregular shape and disruption of cytoplasmic membrane at various regions around the nucleus along with intense vacuolization in most sperm heads (91%; n=103; **Fig. 5A-D**). Cross section of flagellar axoneme of *hsf5*^-/-^mutant spermatozoa showed severe deformity in the arrangement of microtubules. Unlike wild type axoneme, which has a typical ‘9+2’ pattern for microtubule arrangement (30), majority of *hsf5*^-/-^ spermatozoa showed severe structural defects. Some of *hsf5*^-/-^ spermatozoa had a single central microtubule only, while others showed complete lack of central tubules and the rest of the peripheral duplet microtubules were irregularly arranged (85%; n=117; **Fig. 5E-H)**. Longitudinal sections through the flagella also revealed lack of central doublet microtubules and irregular arrangement of central and peripheral microtubules in most of the *hsf5*^-/-^ sperms (**Fig. 5I-J;** see **Fig. 5K** for relative frequency of these defects).

Loss of spermatozoa, reduction in sperm counts and defects in sperm shape and structure in *hsf5*^-/-^ mutants suggest that Hsf5 is very important for spermatogenesis.

### Spermatogenesis in *hsf5*^-/-^ mutant testes appears to be arrested during meiotic division

Increase in the number of primary spermatocytes and loss of spermatozoa in *hsf5*^-/-^ mutants prompted us to examine primary spermatocytes at different stages of meiosis. We performed immunostaining with anti-Hsf5 antibody and anti-Sycp3, a well-known marker for primary spermatocytes (31), to compare meiotic prophase-1 progression between *hsf5*^-/-^ mutant and WT testes.

*hsf5*^-/-^ mutant testes showed a marked increase in the clusters of Sycp3-positive cells, confirming accumulation of primary spermatocytes in comparison to WT, which was suggested by histological analysis **(Fig. 6A&B, Fig 4A-D)**. Detailed analysis of the localization of Sycp3 also allowed us to differentiate between different stages of meiotic prophase in primary spermatocytes, namely, pre-leptotene, leptotene and zygotene/pachytene. In agreement with previously published reports (32), at preleptotene stage, Sycp3 appeared as a small particle at one side of cell **(Fig. 6E)**. At leptotene stage, Sycp3 staining seemed to highlight the bouquet shaped chromosomal arrangement **(Fig. 6F)**, whereas at zygotene/pachytene stage, Sycp3 localization appeared reticulate, staining condensed chromosomes **(Fig. 6G)**. We compared the numbers of cells in these stages between the WT and *hsf5*^-/-^ mutant testes, from randomly selected regions of equal area. *hsf5*^-/-^ mutant testes showed comparatively lower number of cells at preleptotene stage, and significantly higher ones in the subsequent leptotene and zygotene/pachytene stages in comparison to WT testes **(Fig. 6H)**. In addition, the size of the cells at the above stages, was also greater in *hsf5*^-/-^ testes in comparison to that in WT testes **(Supplementary Fig 8)**, which might have resulted from some defects in cell division.

**Fig. 6:** *hsf5*^-/-^ mutant testes showed higher number of primary spermatocytes and apoptotic cells compared to WT. A: WT testis stained with Sycp3. B: *hsf5*^-/-^ mutant testis stained with Sycp3 showing higher number of cells stained by the primary spermatocyte marker. C-D: TUNEL staining of the wild type (C) and *hsf5*^-/-^ (D) testis showing high numbers of apoptotic spermatocytes and spermatozoa in the latter. E: Spermatocytes at preleptotene stage of meiotic prophase 1 where Sycp3 appear as small particle at one side of cell. F: Spermatocytes at leptotene stage where Sycp3 has stained the bouquet shaped chromosomal arrangement. G: Spermatocytes at zygotene/pachytene stage where Sycp3 staining is in reticulate manner. H: The relative proportion of the above three stages quantified from Sycp3-stained WT and *hsf5*^-/-^ mutant testis (n=3 each). Cell counts at the preleptotene stage were significantly lower in *hsf5*^-/-^ mutant, whereas the leptotene and zygotene/pachytene stages were highly represented in the *hsf5*^-/-^ mutant in comparison to WT.

The reduction in the number of post-meiotic cells and their aberrant shape, suggested by the histology analysis, could have resulted from programmed cell death. Indeed, fluorescence-based TUNNEL assay showed significantly higher number of apoptotic cells (both spermatocytes and spermatozoa) in mutants while only a few such cells were observed in WT testis **(Fig. 6C-D)**.

Accumulation of the cells during meiotic leptotene and zygotene/pachytene stages with loss in the cells in post-meiotic stages, suggests that Hsf5 is required for proper progression during meiotic prophase-I in spermatogenesis.

### Hsf5 has a potentially important role in regulating cell cycle and apoptosis

Comparative transcriptome analysis of adult testes from the *hsf5*^-/-^ mutant and wild type individuals revealed that 28% (3,592/12,772) of the genes tested were differentially expressed. Of these, 2,804 were up-regulated and 788 were down-regulated in the mutant with a *p*-value <0.05 **(Supplementary Table 1E)**. A principal component analysis (PCA) of the sequencing data showed that the transcriptome of the *hsf5*^-/-^ zebrafish gonad is significantly different from that of WT **(Fig. 7A)**. The expression of a few *hsp* genes, such as *hsp90ab1*, *hsp70*, *hspa4a*, *hspa8*, *hspa14* and some members of the *hsp40* family, showed marginal up-regulation in mutants, whereas *hsp40c6* and *hspb8* were the only members with slightly down-regulated transcript levels **(Supplementary Table 1E)**.

**Fig. 7:** Transcriptome of the *hsf5*^-/-^ zebrafish gonad is significantly different from that of WT. A: PCA plot demonstrating the clustering of mutants further away from WT. B: Analysis of Hsf5 CHIPseq peak showed a wide distribution with significant number of peaks in intergenic region. C: Functional classification of Hsf5 peaks revealed the mapping of majority of peaks to protein coding genes. D: The intersect between RNAseq DEGs and CHIPseq gene sets showed that a significant number of DE genes were also hsf5 targets. E: KEGG pathway analysis of gene intersect indicated the involvement of DNA repair pathways, meiosis and apoptosis in spermatogenesis.

GO analysis of differentially expressed genes showed enrichment of genes for the following GO terms: ‘Regulation of cell cycle’, ‘Regulation of cell death’, ‘Chromatin organization’, ‘Sister chromatid segregation’, ‘Double strand break repair’, ‘Microtubule based movement’, ‘Intracellular transport’ and ‘ATP binding’, suggesting Hsf5 may be an important regulator of cell cycle during spermatogenesis **(Supplementary Table 1F)**.

Hsf5 loss-of-function resulted in misregulation of 13-38% genes categorized in multiple DNA-repair pathways, including ‘Mismatch repair’, ‘Nucleotide excision repair’, ‘Base excision repair’ and ‘Non-homologous end-joining’ pathways. In addition, nearly 30% of ‘Cell cycle’ pathway genes **(Supplementary Table 1G)**, including cyclins, which play an important role in meiosis (cyclins A1, B1, B3, D2 and E2; for review see: (33, 34)), were also differentially expressed. HSFs are known to contain DBD which, albeit their nuclear localization is observed upon stimulus. A high-resolution imaging showed Hsf5 localization in the nucleus as small speckles in wild type cells **(Fig. 3G, H)**. Therefore, we examined DNA binding of Hsf5 using ChIP-Seq. The results revealed a total of 7,651 peaks with *p*-value <0.05 and >2.5-fold enrichment of peak height in two independent biological replicates compared to input DNA control **(Supplementary Table 1H)**. Nearly 60% of Hsf5 peaks were in the genic regions, mapping to 2,014 genes in the gene bodies, only very few of them were in the promoter region **(Fig. 7B)**. These 2,014 genes were used for investigating the potential role of Hsf5 in the process of spermatogenesis by intersecting with DE genes between *hsf5*^-/-^ *vs*. WT. Many of these Hsf5-bound genes were differentially expressed, among these Hsf5 enrichment was more than 4-fold for 609 DETs **(Fig. 7D; Supplementary Table 1I)**. Of these, 489 (80.3%) were up-regulated and the remaining 120 (19.7%) were down-regulated in *hsf5*^-/-^ mutant males revealing a tight correlation between the genes Hsf5 bound as well as up-regulated upon Hsf5 depletion, suggesting that Hsf5 may function primarily as a transcriptional repressor.

KEGG analysis of Hsf5-bound, differentially expressed genes (*p*-value <0.05 and 2.5-fold or higher enrichment of peak) revealed that a substantial number of these genes were categorized in pathways ‘Cell cycle’, ‘Meiosis’, ‘Apoptosis’, ‘DNA repair’ and ‘Wnt signaling’ **(Fig. 7E; SupplementaryTable 1J** for more details). This suggests that Hsf5 directly regulates the genes required for proper cell division and also explains the meiotic progression failure in the *hsf5*^-/-^mutants.

## Discussion

### Hsf5, a new, conserved heat shock transcription factor is essential for spermatogenesis in zebrafish

Here, we report identification and functional characterization of a novel heat shock factor, Hsf5 from zebrafish. Based on the presence of Hsf5 homologs in more than 30 species (data not shown), including several mammals, chicken and Xenopus, Hsf5 appears to be conserved across the whole vertebrate clade. Our study describes the functional analysis of this new member of Hsf family, in a vertebrate system.

A remarkable feature about Hsf5 seems to be its exclusive function for spermatogenesis in zebrafish, indicated by its expression pattern and detailed analysis of spermatogenesis in *hsf* mutants. Hsf5 expression patterns seem to be conserved among humans and rats too, suggesting its similar function in other mammals (16).

Other orthologues in mouse, *Hsf1* and *Hsf2*, are also transiently expressed in spermatocytes and spermatids (35, 36). Male *Hsf2*-null mice developed smaller testes with abnormal seminiferous tubules resulting in reduced fertility (14). A double knockout of both *Hsf1* and *Hsf2* in male mice resulted in a more severe gonadal phenotype that included infertility due to abnormal sperm shape and reduced sperm numbers (35, 37), similar to what we observed in our *hsf5*^-/-^zebrafish males, suggesting Hsfs play an important role in spermatogenesis.

### Loss of Hsf5 disrupts meiotic progression in spermatocytes

Hsf5 protein localization was predominantly observed in the primary spermatocytes, similarly to Sycp3 ((31); **(Fig. 6A-B)**. The Hsf5 signal was strongest at the cytoplasmic bridges, which interconnect the spermatocytes. In mice and *Drosophila*, these structures were suggested to function as channels to transport gene products (38).

The observed accumulation of primary spermatocytes in the leptotene and zygotene/pachytene stages in zebrafish, strongly suggests that Hsf5 plays a vital role in the progression of meiotic prophase 1. Accumulation might occur from a delay in the cell cycle due to defects, whereas lower number of cells in preleptotene stage in *hsf5*^-/-^ mutant might be the result of a feedback mechanism, where accumulated cells in later states send a signal that inhibits the entry of mature spermatogonia into meiotic stage. Indeed, misregulation of some genes likely explains our observation. For example, follicle stimulating hormone (Fsh) supports entry into meiosis and survival via the intrinsic & the extrinsic apoptotic pathways in rodents (39), the expression level of its receptor (*fshr*) showed a two-fold down-regulation in the *hsf5*^-/-^ mutants. One of the genes with significantly down-regulated expression level in the *hsf5*^-/-^ zebrafish testis is mediator complex subunit 1 *(med1)*. In mouse, MED1 is required for meiotic progression, as Med1 knockout male mice showed reduced number of testicular cells at preleptotene stage and accumulation of spermatocytes at pachytene stage (40). The above results validate our observations and strengthen the role of Hsf5 in proper meiotic progression.

### The expression level of several genes associated with cell cycle regulation or DNA repair is altered in hsf5^-/-^ mutant testes

We observed that several genes regulating cell cycle, apoptosis and DNA damage repair, were misregulated in *hsf5*^-/-^ mutants suggesting that Hsf5 is an upstream regulator of meiosis. Importantly, several cyclin genes such *as cyclin A1*, *cyclin D2*, *cyclin Y and cyclin K* that regulate meiosis showed down-regulated expression in *hsf5*^-/-^ mutants. Cyclin A1 *(ccna1)*, is most abundantly expressed in spermatocytes in the mouse and its deficiency causes meiotic arrest at late meiotic prophase and undergo apoptosis (41, 42). In mice, down-regulation of *cyclin B3 (ccnb3)* expression is necessary at the zygotene-pachytene transition for the normal progression of spermatogenesis (43). The expression of cyclin B3 was up-regulated in our mutant zebrafish males leading to abnormal spermatogenesis. E-type and D-type cyclins also play an important role in prophase 1 of meiosis in mice. Down-regulated expression of cyclins involved in spermatogenesis provides a glimpse of molecular details that underlie the meiotic progression failure in *hsf5*^-/-^ mutants.

In addition, we also noticed a down-regulation of 25 members of PIM kinases, **(Supplementary Table 1E)**. These proteins belong to CAMK group of kinases, which regulate cell cycle and growth. Knockdown or inhibition of PIM-1 was correlated to increased apoptosis, induction of heat-shock family proteins (44) and inhibition of nonhomologous DNA-end-joining (NHEJ) repair activity in humans.

Hsf5 loss-of-function in zebrafish caused misregulation of up to a third of genes categorized in ‘Mismatch repair’, ‘Nucleotide excision repair’ and ‘Base excision repair’ pathways. DNA repair is a basic process essential for the maintenance of genomic integrity by playing a critical role in mitosis and meiotic recombination and chromosome pairing (45).

The cAMP responsive element modulator (*crema*) is down-regulated in *hsf5*^-/-^mutant testes. Loss of function of its mouse orthologue CREM, leads to severe impairment of spermatogenesis and germ cell apoptosis. CREM is regulated by germ cell-specific kinesin, Kif17b. Thirteen members from *kif* family, including *kif17*, were differentially expressed in our study suggesting that Crema might also play a role in zebrafish spermatogenesis.

The expression of several members of the regulatory factor X *(rfx)* family of transcription factors, such as *rfx7*, *rfx1a* and *rfxap*, was also up-regulated in *hsf5*^-/-^males. The RFX2 transcription factor is a master regulator of genes required for the haploid phase in mice and Rfx2^-/-^ mice show complete male sterility (46). RFXs regulate intraflagellar transport (IFT) genes and RFX1 is reported to be essential for later stages of spermatogenesis (47). In our data, we also noticed enrichment of genes for GO terms ‘Sister chromatid segregation’ and ‘Double strand repair’ indicating the potential presence of a compensatory mechanism in *hsf5*^-/-^ males. A similar mechanism has also been proposed earlier for human spermatogenesis (48).

### Down-regulation of important cytoskeleton and motor proteins explain the defects with sperm mobility and short tails in hsf5^-/-^ mutant males

Disruption of proper microtubules arrangement in the sperm axoneme is likely explained by misregulation of cytoskeletal and motor proteins. Axonemal dyneins are considered “arms” that are attached to each of the nine doublet microtubules (49). Our data showed reduced expression of *dnai1.2* in the mutant, whereas *dnai2a* and six members of the *dnah* gene family *(dnah1*, *dnah5*, *dnah7l*, *dnah9*, *dnah9l* and *dnah10)* all showed up-regulated expression levels compared *to* WT. Misregulation of these genes have been reported in the sperm motility related disorders. In ciliary dyskinesia patients, abnormal assembly of DNAH7 was identified previously (50). Similarly, defects in human DNAI1 resulted in primary ciliary dyskinesia type 1 (CILD1) with abnormalities in the sperm flagellum and reduced fertility (51). Misregulation of cytoskeletal coding genes, such as *map1sb*, *saxo2*, *msat4* and *mapre2*, likely explains abnormal microtubule arrangement of axonemes in 85% of *hsf5*^-/-^ mutant zebrafish spermatozoa. In addition, we also observed down-regulated expression of many tubulin genes, such as *tuba2*, *tubcd*, *atat1*, *ttll1*, and *ttll3*, in *hsf5*^-/-^ mutant males.

Intraflagellar transport (*ift*) genes are essential for the bidirectional movement of substances within cilia and flagella as well as maintenance and assembly of these structures (52, 53). Over-expression of three *ift* genes *(ift20*, *ift80* and *ift172)* might have contributed to the abnormal flagella formation and sterility of *hsf5*^-/-^male fish. Spag6, an axoneme central apparatus protein, is essential for the function of ependymal cell cilia, sperm flagella and axoneme orientation. Many *Spag6*-deficient mice showed sterility because of sperm motility defects (54). In our study, *spag6* expression level was reduced in *hsf5*^-/-^ mutants, whereas *spag1b*, *spag7*, *spag9* and *spag16*, were all expressed at a higher level compared to WT.

Dynein axonemal assembly factor (Dnaaf) proteins are important for assembling axonemal dynein and stability of cilia and flagella (55-57). In human patients lacking DNAAF2, both outer and inner dynein arms were partially or completely absent and sperm flagella were immotile (58). *hsf5*^-/-^ zebrafish males also showed lower *dnaaf2* transcript levels than WT. Misregulation of these genes coding for cytoskeleton and motor proteins points to potential importance of Hsf5 in the control of genes associated with sperm mobility.

Hsf expression is mostly reported in the cytoplasm, the localization of Hsf1 and Hsf2 to stress granules in nuclei upon heat shock has been reported in humans (59, 60). Since Hsf5 loss of function did not result in major misregulation of *hsp* genes, it does not seem to function in the heat shock response, however, its nuclear accumulation as speckles, in primary spermatocytes, was visible. Remarkably, many genes with significant Hsf5 binding were misregulated in the hsf5 mutants, suggesting its role in transcriptional regulation of those important genes. Hsf5 peaks in the gene bodies suggest, that it may not act as canonical transcriptional regulator, which bind to the promoter regions. Future analysis of epigenomic status of DEGs in Hsf5 mutants and Hsf5 interacting partners may reveal more about the mechanistic details of Hsf5 function.

## Materials and Methods

### Fish husbandry

This study and all procedures were approved by Temasek Life Sciences Laboratory Institutional Animal Care and Use Committee (approval ID: TLL(F)-10-002) for experiments carried out at Temasek Life Sciences Laboratory (License for Animal Research Facility No. VR016) and Örebro University by Linköpings djurförsöksetiska nämnd (Linbköpings Animal Care and Use Committee, Approval ID: 32-10) for experiments carried out at Örebro University (License for Animal Research Facility: No. 5.2.18-2863/13).

Zebrafish (*Danio rerio*, AB strain) were raised, maintained, and crossed according to the standard protocol (61). Samples of different stages of gonad development were collected using Tg(ddx4:ddx4-egfp) (formerly known as Tg(vas:egfp) transgenic zebrafish line (a generous gift from Dr. Lisbeth Oslen, SARS, Bergen, Norway). Fish were reared in AHAB recirculation systems (Aquatic Habitats, Apopka, FL, USA) at ambient temperature (26-28°C).

### Identification of hsf5 and its two isoforms

The expressed sequence tag (EST) with testis-enhanced expression was identified from our full-length, normalized adult testis cDNA library (23, 25). A BLASTn search with this sequence as a bait yielded seven other testis-derived ESTs from GenBank. The consensus sequence (1,581bp) was confirmed by RT-PCR (for the complete list of genes analyzed in the study and primers used for their amplification please see **(Supplementary Table 1B)**. To obtain the complete *hsf5* cDNA sequence, rapid amplifications of cDNA ends (RACE) were performed using RLM-RACE kit (Ambion). The amplified products were cloned into pGEM-T easy vector (Promega) and used for sequencing.

The coding region of zebrafish *hsf5* cDNA was amplified from various organs and different developmental stages using primers annealing to the ends of the cDNA and Qiagen 1 step RT–PCR kit. The products from PCR reactions were cloned into pGEM-T easy vector (Promega) and sequenced for verification. The expression of *hsf5* transcript variants was analyzed by RT-PCR using a primer pair (hsf5 FL) that amplified both *hsf5_tv1* and *hsf5_tv2*. The following PCR cycling conditions were used: initial denaturation step 95°C for 1 min, then 95°C for 30s, 60°C for 30s, 72°C for 30s; 36 cycles. *eef1a1l1* was used as a reference gene. Fully sequenced PCR products were compared to determine the difference between the two variants.

### Bioinformatics and phylogenetic analysis of Hsf5

The Hsf5 protein was analyzed using the following softwares: Conserved Domain Database (CDD), SMART (both available at: http://expasy.org/). The full-length amino acid sequence of Hsf orthologs were retrieved from GenBank **(Supplementary Table 1C)**. The sequences were aligned by the CLUSTAL OMEGA software (62). Estimation of molecular phylogeny was carried out by the neighbor-joining method with Poisson correction model as implemented in MEGA (Version 5) (63). Confidence in the phylogeny was assessed by bootstrap re-sampling of the data (1000) (64).

### Sample collection and isolation of nucleic acids

For the analysis of early gene expression, RNA was extracted from pooled zebrafish embryos (30 individuals/pool) collected at 3 days post-fertilization (dpf). Snap-frozen samples were stored at −80°C until use.

In order to investigate the expression of *hsf5* during gonad development, samples from the isolated gonads (20, 25, 30, 35 and 40 dpf) were collected from a total of six Tg(ddx4:ddx4-egfp) individuals for each time point. Previously, our lab showed that those transgenic individuals that did not seem to show visually detectable Egfp signal during their early gonadal development (20-24 dpf) became exclusively males (65). Therefore, we considered such individuals as presumptive males (four samples), whereas individuals with strong Egfp signal were considered as presumptive females (four samples). Individuals were sorted into these two categories as described earlier (65).

For spatial analysis of *hsf5* expression profiles in adult zebrafish, samples from nine different organs (testis, ovary, kidney, liver, brain, gut, gill, skin and eye) were collected from three individuals. For heat shock experiment, two groups were formed from adult male siblings. The first group was heat-treated for one hour at 37°C, whereas the second group was kept at ambient temperature (26-28°C). Testis samples were collected from three individuals in each group. Total RNAs were extracted using either RNeasy RNA extraction kit (Qiagen) or TRIzol-LS reagent (Life Technologies) according to the manufacturer’s instructions. RNA isolates were quantified using an ND-1000 spectrophotometer (NanoDrop). First-strand cDNA was synthesized under standard conditions using iScript cDNA synthesis kit (Bio-Rad) for the analysis of embryonic expression, whereas the Superscript First-strand Synthesis System (Invitrogen) using an oligo (dT)_15_ primer (Roche) was used for adult organ samples.

### Gene expression analyses

The expression analyses of *hsf5* during early embryonic development, gonad development and in adult tissues were performed by using real-time quantitative PCR (qPCR) using the iCyler iQ Real-time Detection system and SYBR Green chemistry (Bio-Rad). For the quantification of the transcripts, a standard curve with 10-fold serial dilution of testis cDNA was used. The 15 μl qPCR mixture contained 7.5μl of 2× iQ™ EVA Green Supermix, 0.5 μl (5 mM) of each primer (these primers amplify both isoforms) and 1 μl of cDNA. The following conditions were used: denaturation at 95 °C for 3 min, 40 cycles of denaturation at 95 °C for 30s, annealing at 60 °C for 30s and extension at 72 °C for 20s, followed by melt curve analysis (95 °C for 2 min and decrease by 0.1 °C at every 10s). Samples were assayed in triplicate and each experiment had at least three biological replicates. To normalize the expression, several housekeeping genes (including *actb1, rpl13*, and *eef1a1l1)* were assayed in order to identify genes with stable expression among the organs sampled and across different developmental stages. We found that both *rpl13* and *eef1a1l1* were suitable for the analysis of early developmental stages and adult zebrafish tissues, whereas for the study of developing gonads, *rpl13* worked well. In the present study, we used *eef1a1l1* for early developmental stages and adult tissues, whereas *rpl13* was used as a reference for the developing gonads.

The efficiency of each reaction was calculated using PCR miner (66). The relative gene expression level was determined using the delta-CT method of normalizing gene expression by subtracting the average reference Ct value from that of the average target and is presented as a log2 of relative quantity (RQ) of target gene.

### Immunohistochemistry

For immunohistochemical (IHC) analyses, testes and ovaries from adult zebrafish (90 dpf) were isolated and fixed in 4% paraformaldehyde in PBS (pH 7.4; Sigma) for 2 hours (hrs) at room temperature (RT) and washed three times with PBS and equilibrated in graded sucrose series. Samples were frozen in tissue freezing media (Leica Biosystems), and 5-7 μm sections were cut using a cryotome (Leica Biosystems). Nonspecific protein binding sites were blocked by 30 mins incubation in PBS-based blocking buffer containing 3 % BSA (Sigma) and 0.2 % Triton X. The sections were then incubated either with anti-Hsf5 or anti-Sycp3 antibody for 16 hrs at 4 °C at 1:1000 and 1:400 dilutions, respectively, in PBS containing 1 % BSA and 0.1 % Triton X. Alexa Flour 488 anti-rabbit (Invitrogen) at 1:1000 dilutions were used as a secondary antibody to incubate for 2 hrs at RT followed by DAPI (Calbiochem) staining for 5 mins. The image was captured using an SP8 gSTED Confocal Laser Scanning Biological Microscope (Leica). Three randomly chosen images stained with Sycp3 from three biological replicates were used for counting different cell types in meiotic prophase stages. To determine the meiotic stage of the primary spermatocytes we took advantage of Sycp3 antibody staining. The nuclei were stained by DAPI (red).

### Generation and screening of mutants

Custom-designed guide RNA (gRNA) targeting the DNA Binding Domain at second exon of *hsf5* and recombinant Cas9 protein (*Streptococcus pyogenes*) were ordered from ToolGen Inc. gRNA was designed using CRISPR design tool (http://crispr.mit.edu/) and off-target analysis using RGEN Tools by Seoul University (http://www.rgenome.net/cas-offinder). Based on the above analysis result, a gRNA with no off-target effect was synthesized (**Supplementary Fig 9**) for the position and sequence of the gRNA targeting site).

gRNA and Cas9 protein were co-injected into 350 one-cell stage zebrafish embryos in three separate experiments. Each embryo was injected with 2 nl of solution containing 12.5 ng/ml of sgRNA and 300 ng/ml of Cas9 protein. Injected embryos were grown to 60 dpf for fin clipping. A total of 57 F0 founder individuals were screened by fluorescent genotyping and 16 of them by T7E1 assay (for typical results see **Supplementary Fig 10**) as well.

For screening, genomic DNA from the tail fin clip of three mpf (months post-fertilization)-old CRISPR/Cas9 injected zebrafish was isolated using genomic DNA extraction kit (Qiagen) following the manufacturer’s protocol. The genomic region surrounding the CRISPR target site was PCR amplified using primer pair ‘hsf5a’ and cloned into pGEMT Easy vector (Promega). Positive clones were selected and the extracted plasmids were sequenced by Sanger method. For T7E1 assay, 200 ng PCR product per individual was subjected to a re-annealing process to enable heteroduplex formation: 95 °C for 10 min, 95 °C to 85 °C ramping at 2 °C/s, 85 °C to 25°C at 0.1 °C/s and holding at 25 °C for 1 min. After re-annealing, 5 units of T7 Endonuclease I (NEB) was added to the PCR products following the manufacturer’s recommended protocol, and they were separated on 2% agarose gels in TBE buffer. Relative proportion of mutation was estimated by the formula of Indel % = 100 x (1- √(1-fcut)) as described in (67).

Ninety per cent of the tested founders contained mutations in the 2^nd^ exon of the *hsf5* gene (for typical results see **Supplementary Fig 6)**, the estimated percentage of mutation ranged from 17% to 54%. That exon was sequenced from sixteen selected F0 mutants to assess the number and type of mutations per individual and the results have shown the presence of several different insertions and/or deletions in each of these founder individuals **(Supplementary Fig 11)**.

### Propagation and genotyping of mutants

F0 mosaic founders were raised to sexual maturity and crossed with wild-type (WT) partners to generate F1. The resulting embryos were grown to three mpf, genomic DNA was isolated from tail fin biopsies and subjected to genotyping to identify the mutants. Heterozygous mutants from F1 were crossed to generate F2 for subsequent analyses.

For genotyping, the forward primer of ‘hsf5b’ primer pair was labeled with FAM fluorescent dye on the 5’ end and PCR was performed in a 25 ul volume to amplify a 267 bp fragment spanning the targeted region of the *hsf5* locus. PCR products were mixed with internal size standard, GeneScan 500 Rox (Applied Biosystems) and subjected to capillary electrophoresis using 3730xl DNA analyzer (ABI, Foster City, CA, USA). The genotypes of mutants were analyzed using Genemapper software (Version 5.0).

For fertility tests, twelve WT male and twelve WT female zebrafish were selected for pairwise crossing. The total number of eggs produced and survival rate at 24 hpf was recorded for three to six rounds of breeding. After that, six and three WT females with consistent production of good quality eggs were selected to be paired with *hsf5*^-/-^ and *hsf5*^+/-^ male partners, respectively. Eight WT males were paired with *hsf5*^-/-^ females and the transparent eggs from all were collected separately after 3 hpf to determine the survival rate at 24 hpf.

### Histological and SEM analysis of gonads

Male zebrafish were euthanized with tricaine methane sulfonate and their testicular tissues were fixed in 4% formaldehyde at 4°C overnight. After dehydration, samples were embedded in plastic resin (Leica). Serial cross-sections of 2 μm were cut by microtome (Leica), dried on slides at 42° C overnight, stained with haematoxylin and eosin, and then mounted in Permount (Fisher-Brand) and imaged with phase contrast microscope with a 63x and 100x oil objective lens. Four WT and *hsf5*^-/-^ testis were used for morphometric analysis of germ cell types. Ten sections were chosen randomly for each sample and scored using high resolution light microscope (Leica) at a magnification of ×100. Staging of germ cells was performed as described previously (68), using collection of images explaining the histology and toxicological pathology of the zebrafish (69) as a reference. ImageJ thresholding was used to count the number of germ cells. Images with sectioning or staining artefacts were not selected for cell counting.

For Scanning Electron Microscopy (SEM), three wild type and mutant sperm specimens each were fixed using 2.5% glutaraldehyde, dehydrated in ascending graded ethanol (30%, 50%, 70%, 90% and 96% for 15 minutes in each) then dropped onto a round glass slide and dried using a silica gel drier. After the surface was treated with electric conduction, the specimens were observed under a Jeol JSM-6360LV scanning electron microscope.

For Transmission Electron Microscopy (TEM), samples (n=3) were fixed with 2.5% glutaraldehyde in 0.1 M phosphate buffer (pH 7.2), washed 3X in 0.1 M phosphate buffer (pH 7.2) for 15 minutes each dehydrated in graded series of alcohol (in water) for 15 minutes each, dried at critical point, mounted on specimen stub with silver paste, sputter coated with gold and imaged under a Jeol JEM-1230 Transmission Electron Microscope (Jeol, USA).

### Sperm quantification and TUNEL assay

Sperm counts were determined by hemocytometry. Sperm suspension stock was generated by crushing the full testes of each individual in Fifty Microliter Hank’s balanced salt solution. This stock from four WT and mutant individuals each was diluted 5000 times, and was loaded to fill the haemocytometer counting chamber. Sperm numbers were counted using light microscope at 20X magnification. Spermatozoa in four corner squares were counted from four replicates of the same individual sample and used for calculating the concentration.

TUNEL staining was performed on mutant and wildtype testis fixed in 4% paraformaldehyde, embedded in paraffin. Cross sections of 5 μm were rehydrated and a commercial *in situ* cell-death detection kit (Roche diagnostics, Germany) was used for labelling the slides. Nonspecific protein binding sites were blocked by 30 mins incubation in blocking buffer containing 3 % BSA (Sigma) in 0.2 % Triton X in PBS. The sections were then incubated with antifluorescein antibody (Invitrogen) for 16 hrs at 4 °C at 1:400 dilution in PBS containing 1 % BSA and 0.1 % Triton X. Alexa Flour 488 anti-rabbit (Invitrogen) at 1:1000 dilutions were used as a secondary antibody for 2 hrs at RT followed by DAPI (Calbiochem) staining for 5 mins. The image was captured using SP8 gSTED Confocal Laser Scanning Biological Microscope (Leica).

### Transcriptome analysis and CHIP sequencing

RNA was extracted from the intact testes of adult *hsf5*^-/-^ and WT siblings (at five mpf of age) using an Ambion RNAqueous-micro kit (Life Technologies). Sequencing libraries were constructed using TruSeq RNA Library Prep Kit v2 (Illumina) following the manufacturer’s instructions and sequenced using Illumina Nextseq 500. The 150 bp paired-end reads generated were pre-processed using Cutadapt v.1.18 (70) and Trimmomatic v0.36 (71) to remove low quality reads. Mapping was performed on GRCz10 genome assembly using STAR v2.5.3a (72). Differentially expressed genes were identified using Partek Flow software (Partek Inc). Genes with FDR values of less than 0.05 and average coverage of at least 5 were classified by GO (Gene Ontology) and KEGG (Kyoto Encyclopedia of Genes and Genomes) pathway analysis. Unclassified genes were annotated by referring to their orthologous genes in mammals.

An antibody was raised against the zebrafish Hsf5 protein by Agrisear AB (Vännäs, Sweden) against the C-terminal region (amino acids 407-420), which is not conserved among the Hsf5 orthologs in vertebrates. The specificity of the antibody to recognize native Hsf5 protein was demonstrated through Western blot and immunohistochemistry on zebrafish gonads. Chromatin Immunoprecipitation (ChIP) was performed using MAGnify™ Chromatin Immunoprecipitation System (Invitrogen) following the manufacturer’s instructions. Library preparation was performed on 50 ng CHIP DNA using the Illumina ChIP-seq sample preparation kit (Illumina, USA) according to the manufacturer’s protocol. The ChIP DNA library was sequenced on the Nextseq 500 (Illumina, USA) with paired end, 150 bp reads.

## Statistical Analysis

For qPCR data analysis, mean ± standard deviation was calculated. Statistical differences in relative mRNA expression between experimental groups were assessed by 2 tailed Student’s *t*-test. Differences were considered statistically significant at *p*<0.05. *, p <0.05; **, p < 0.01; ***, p < 0.001.

We used Kolmogorov-Smirnov test to compare the significance in the difference between the cell number and area at meiotic prophase-I.

Data: The transcriptomic and ChIP-seq data in the current study is submitted to

NCBI SRA, with accession number <u>SRP124146</u>

## Author contributions

Conceived the study: LO, JMS, MHS,

Designed the experiments: JMS, AA, PEO, LO

Generated the mutants: JMS

Performed the experiments and/or analyzed the data: JMS, MHS, AP, WCL, NMT

Discussed and interpreted the data: JMS, MSH, AP, WCL, NMT, AA, PEO, LO

Wrote and corrected the manuscript: JMS, AA, PEO, LO

## Acknowledgements

This research was supported by the National Research Foundation, Prime Minister’s Office, Singapore under its Competitive Research Programme (Award No: NRF-CRP7-2010-001) and by internal research grants from Temasek Life Sciences Laboratory (for LO), by fellowships from Temasek Life Sciences Laboratory and the Department of Biological Sciences of the National University of Singapore (for MSH) as well as by the Swedish Research Council, Knowledge Foundation, Sweden and Örebro University (for PEO).

The authors thank the following colleagues for their helpful technical advices: Norman Teo on CHIP experiment, Kellee Siegfried on histology, Toshihiro Kawasaki on the IHC procedure, Haiwei Song, Bharath SR and Xuhua Tang on analyses of protein domains. Sycp3 antibody was a kind gift from Noriyoshi Sakai.

## Supplementary Information

**Supplementary Fig 2. Hsf5 does not play a role in heat shock response**. A: Validation of Hsf5 antibody in testis showing manifold higher expression compared to ovary. B-C: Transcripts levels of *hsp70* in testis showed upregulation upon heat-shock in adults, whereas *hsf5* levels remained same after heat-shock. Transcript levels were normalized against *eef1a1a* reference gene. Each value represents the mean of n=3 +/- SD. * *p*<0.05; **p<0.01:*** p<0.001, Student t-test.

**Supplementary Fig 3. *hsf5* expression was low during early embryonic development and among the organs tested testis showed the highest expression**. A: Low expression levels from of *hsf5* detected from zygote to juvenile (21dpf) by q-PCR. **B**: Testis showed highest expression level compared to other organs (*eef1a1l1* was used as a reference gene; n=3). Transcript levels were normalized against *eef1a1a* reference gene. Each value represents the mean of n=3 +/- SD. * *p*<0.05; **p<0.01; *** p<0.001, Student t-test.

**Supplementary Fig 4. *hsf5* is a late testis marker in zebrafish**. The expression level of *amh* (**A**), *nr5a1a* (**B**) *cyp19a1a* (C) and *hsf5* (D) in both sexes during the gonad differentiation period (20-40 dpf) in zebrafish. Zebrafish transgenic to the Tg(ddx4:ddx4-egfp) *zf45* construct were sorted according to their gonadal Egfp signal: no signal – male; strong signal – female. RNA was isolated from isolated gonads (20-40 dpf). Transcript levels were normalized against *rpl13* reference gene. Each value represents the mean of n=3 +/- SD.

**Supplementary Fig 5. Comparison of two *hsf5* transcript variants in zebrafish testis**. A: RT-PCR using hsf5 full length (Hsf5 FL) primer detected two transcript variants of *hsf5* in testis, whereas the ovary showed only the longer transcript variant. B: Comparative sequence analysis of the two transcript variants, *hsf5_tv1* and *hsf5_tv2*, showed that the testis-specific one *(hsf5_tv2)* was shorter due to the absence of the third exon. Red color arrows indicate the binding sites of primers used for PCR amplification

**Supplementary Fig 6. Efficient germline transmission was observed in F1 generation**. Cross between mosaic F0 males and females with WT yielded A) 21 to 69% germline transmission to F1. B-D: Fluorescent genotypes of B) Wild Type; C) *hsf5*^-/+^ and D) *hsf5*^-/-^

**Supplementary Fig 7. Hsf5 loss of function resulted in male infertility**. Embryo phenotypes for wild type (A,B) and *hsf5*^-/-^(C,D) at 6 (left) and 24 hours post-fertilization (hpf; right).

**Supplementary Fig 8. The surface area of mutant cells was higher than that of WT at meiotic prophase stages**. A-C: Pre-leptotene, leptotene and zygotene/pachytene stages showed significantly higher surface area in *hsf5*^-/-^mutant compared to WT. WT and *hsf5*^-/-^ mutant testis stained with Sycp3 were used for surface area analysis (n=3).

**Supplementary Fig 9. The position of the gRNA targeting site in the second exon of the zebrafish *hsf5* locus**. The target site sequence is indicated in red, whereas the protospacer adjacent motif (PAM) is shown in green. Primer binding sites are in blue.

**Supplementary Fig 10. T7 E1 Assay on F0 founder A) males and B) females showed different rates of mutation (17-54%)**. Indel % = 100 x (1-√(1-fcut))

**Supplementary Fig 11. Mutations generated by engineered gRNA/Cas9 at *hsf5* site in four injected founder individuals**. The wild type sequence is shown at the top. Target sites are blue and the PAM sequence is red and underlined. Deletions are shown as red dashes and insertions blue highlight. (Labels on the right: +: insertion; –: deletion).

**Supplementary Table 1A:** The complete list of zebrafish genes analyzed in this study

**Supplementary Table 1B:** The complete list of primer sequences used for PCR-amplifications

**Supplementary Table 1C:** Vertebrate Hsf orthologs used to generate the phylogenetic tree in Figure 2.

**Supplementary Table 1D:** Percentage of amino acid identity of Hsf5 protein and functional domain between zebrafish and other species

**Supplementary Table 1E:** The complete list of differentially expressed genes from RNA seq

**Supplementary Table 1F:** GO analysis of differentially expressed genes

**Supplementary Table 1G:** KEGG pathway analysis of differentially expressed genes

**Supplementary Table 1H:** ChIP-seq peaks for Hsf5

**Supplementary Table 1I:** Gene list for RNAseq and CHIP seq intersect (*p*-value <0.05 and 4 fold or higher enrichment of peak)

**Supplementary Table 1J:**. KEGG pathway analysis of genes from RNAseq and CHIPseq intersect.

